# Dendrite Control Theory: A Unifying Framework Bridging Neuronal Morphology And Dendritic Voltage Behavior

**DOI:** 10.64898/2026.11.04.764328

**Authors:** Fereshteh Lagzi, Alexander Bird, Hermann Cuntz, Romain Caze, Felix Rosenow, Albert Gidon, Boris Gutkin, Peter Jedlicka

## Abstract

Control theory and dendritic computation have revolutionized their respective fields, yet they rarely converge. Here, we use control theory as a framework to reveal how dendritic morphology and synaptic input organization constrain structural resilience and energy cost. Treating diverse dendritic trees as connected network nodes, we demonstrate that the degree of structural symmetry dictates easy- versus hard-to-access control states. Notably, control inputs restricted solely to peripheral terminal branches are mathematically sufficient to govern the voltage states of the entire dendrite and soma, granting striking resilience against input loss elsewhere. Conversely, limiting input dimensionality amplifies voltage fluctuations and metabolic costs. These findings expose fundamental biophysical trade-offs between wiring cost, functional flexibility, robustness, and energy efficiency, suggesting interneurons generally exhibit higher resilience while excitatory neurons feature superior controllability. We introduce “Dendrite Control Theory” as a rigorous engineering framework linking neuronal morphology directly to computation and robustness, offering a new neuroscience research paradigm.

## Introduction

In recent decades, it has become clear that dendrites are not passive cables, but complex, dynamic systems with rich local and global activity (Stuart and Spruston, 2015; Poirazi and Papoutsi, 2020; Stingl et al., 2025). Techniques such as dendritic patch-clamp recordings, calcium imaging, and voltage-sensitive dye imaging have revealed a diverse range of local and global dendritic and somatic dynamics (Häusser et al., 2000; Larkum et al., 2022; Stuart and Spruston, 2015; Stuyt et al., 2022; Wong et al., 2026). For example, dendrites exhibit diverse behaviors at different scales, from plateau potentials at the sub-branch level (Antic et al., 2010; Holthoff et al., 2006; Larkum et al., 2022; Major et al., 2008; Schiller and Schiller, 2001; Stuyt et al., 2022), through global behavior such as forward-propagating dendritic spikes (Golding et al., 2002; Gooch et al., 2022; Kim et al., 2015; Larkum et al., 2022), to back-propagation-activated calcium spikes (BACs) that unite dendrite and soma (Fig. 1A). The voltage state across an entire dendrite can influence computations beyond what is seen in the somatic output (Poirazi et al., 2003b; London and Häusser, 2005), and is important for ongoing synaptic plasticity (Graupner and Brunel, 2012; Bittner et al., 2017; Ebner et al., 2019). However, a systematic framework that explains how diverse neuronal morphologies can shape these local and global dynamics is currently missing. Moreover, it is unclear how the morphologies of specific cell types contribute to the functions each cell type serves in the brain.

**Figure 1:**
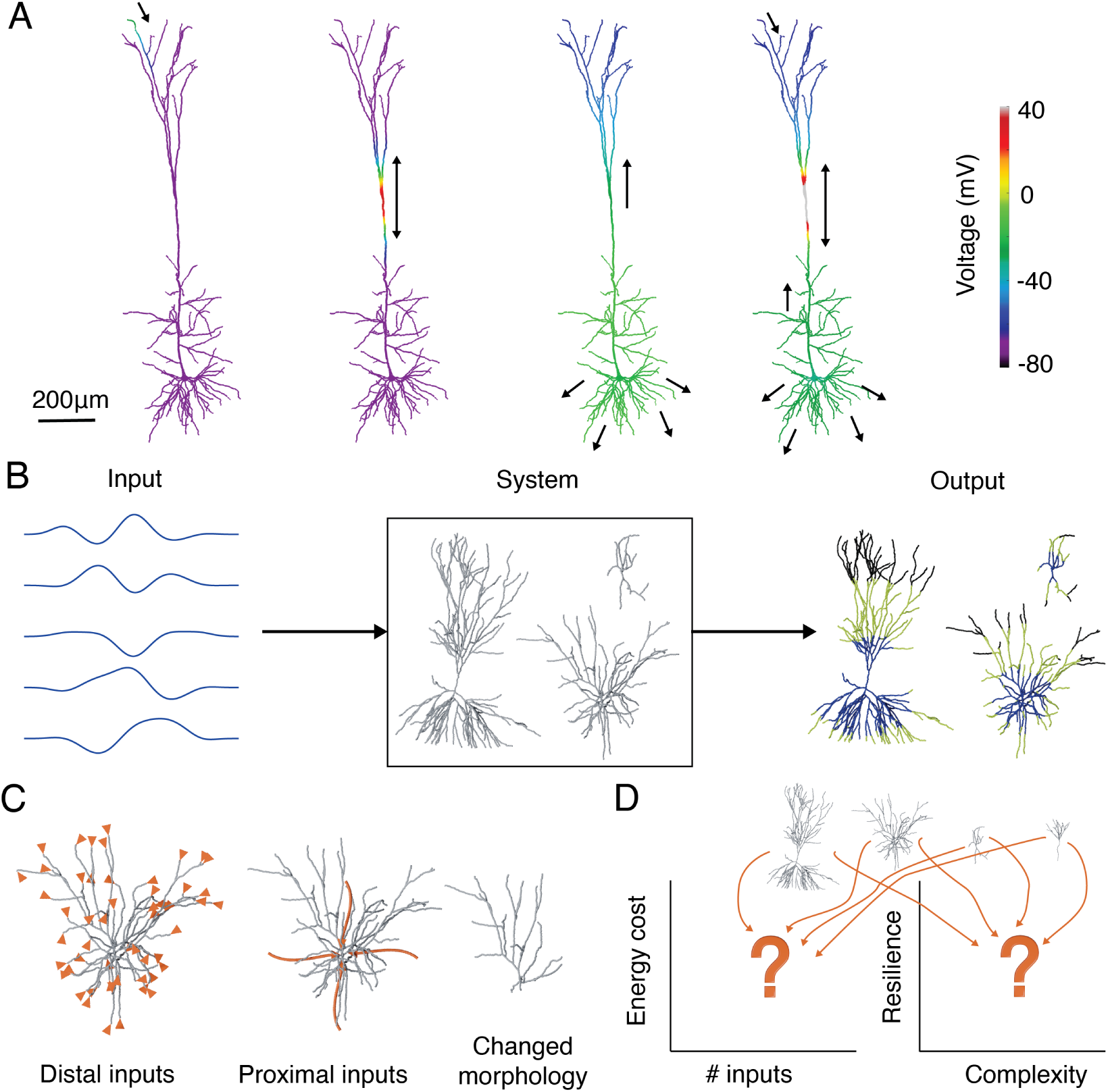
Dendrite control theory. **A** Local and global dendritic states in a layer V pyramidal neuron. Color represents local voltage, and black arrows show the direction of current flow. Left to right: Sub-branch activation, a plateau potential. Global dendritic event, a dendritic calcium spike. Somatic and local dendritic events, a backpropagating action potential after 0.5ms. Global neuronal event, BAC firing after 0.5ms. Morphology from Hay et al. (2011). **B** Control theory framework: What input is necessary for a given system (neuron) to produce a desired output? **C** Changing dendritic inputs and morphology can be viewed as a control mechanism affecting the neuronal state. **D** Could designing control inputs help us identify which cell type is more resilient to structural changes, and which cell type is more energy efficient?

To address these questions, computational studies have modeled dendrites as high-dimensional dynamical systems (Poirazi et al., 2003b; He et al., 2026) and have shed some light on how structure can influence dendritic states. However, a more system-level understanding of the role of morphology in cellular functions, the energy demands to perform dendritic computations, and also the robustness of individual cell types to a wide range of disturbances and morphological restructuring is lacking.

Control theory offers a formal framework to address these challenges. It has been developed and applied across multiple fields to assist with system identification. More precisely, control theory offers principles to design an input signal or shape a dynamical system such that the output serves a desired function (Fig. 1B). This principled design approach can provide useful insight into the nature of the original system.

In this article, we propose that concepts and tools from control theory can help us i) understand how dendritic topology and biophysics shape dendritic dynamics, ii) how neurons can optimally perform specific tasks and computations (Bicknell and Häusser, 2021; London and Häusser, 2005; Segev and London, 2000) within their circuits, iii) how perturbations to dendrites can influence their robustness and if those perturbations can teach us about their functions (Fig. 1C), iv) design optimal stimulation paradigms for achieving desired target states and for probing inherent dendritic voltage (and calcium) dynamics. Hence, we provide a system-level formalism, called Dendrite Control Theory (DCT), and link the properties of dendritic voltage behavior to their morphologies and inputs.

More specifically, using this novel framework, we illustrate how dendritic morphology can influence the voltage states that are easily attainable, finding a trade-off between robustness, complexity, and energy demands (Fig. 1D). Different morphologies imply different trade-offs and, therefore, would be suited to different functional roles within a circuit. DCT provides a new perspective on the structure-function relationship of neurons and interneurons with varied morphologies (Ascoli et al., 2007; Petilla Interneuron Nomenclature Group, 2008). It also explains how morphology could impact the energy required to achieve desired states. Further, we elaborate on the special features of dendrites as dynamical systems that arise from their underlying tree structure. We find that diverse biological phenomena, such as dynamic inhibitory receptor sizes on the distal dendrites (Boivin & Nedivi., 2018; Gidon and Segev, 2012), a wide distribution of synapses across the dendrite (Kasthuri et al., 2015), and the lack of oscillatory dynamics in passive dendrites can be accounted for by DCT.

## Individual neuron conceptualized as a network of interacting nodes

Single neurons often have elaborate dendritic trees with their own complex biophysical and computational properties (Johnston and Narayanan, 2008; London and Häusser, 2005; Stuart and Spruston, 2015). To simplify such complex systems with different morphologies, computational approaches have suggested discretizing dendritic elements into simpler nodes with defined input-output functions and studying interactions between these nodes to better understand the computations performed by neurons.

Nodes in a dendrite can be defined in different ways depending on the scientific question at hand. A node could depend on a stereotypical nonlinear response (Poirazi et al., 2003b), on compartmentalization performed on inputs (Cuntz et al., 2010; Wybo et al., 2019), or on simply sampling at a uniform physical (or electrotonic) rate (Hines, 1993; Rall, 1964; Rall and Rinzel, 1973). In any case, a dendrite (Fig. 2A, left) represented as a set of nodes (Fig. 2A, center) must be coupled together with an appropriate coupling matrix (Fig. 2A, right). The entries in the coupling matrix represent the instantaneous transfer resistance between any pair of nodes. This will be zero for non-adjacent nodes and will otherwise depend on the length, diameter, axial resistance, membrane conductance, and capacitance of the two nodes. A set of dendritic nodes and their coupling matrix can be used together to model a single dendrite as a network (Cuntz et al., 2007; Mittal and Narayanan, 2024).

**Figure 2:**
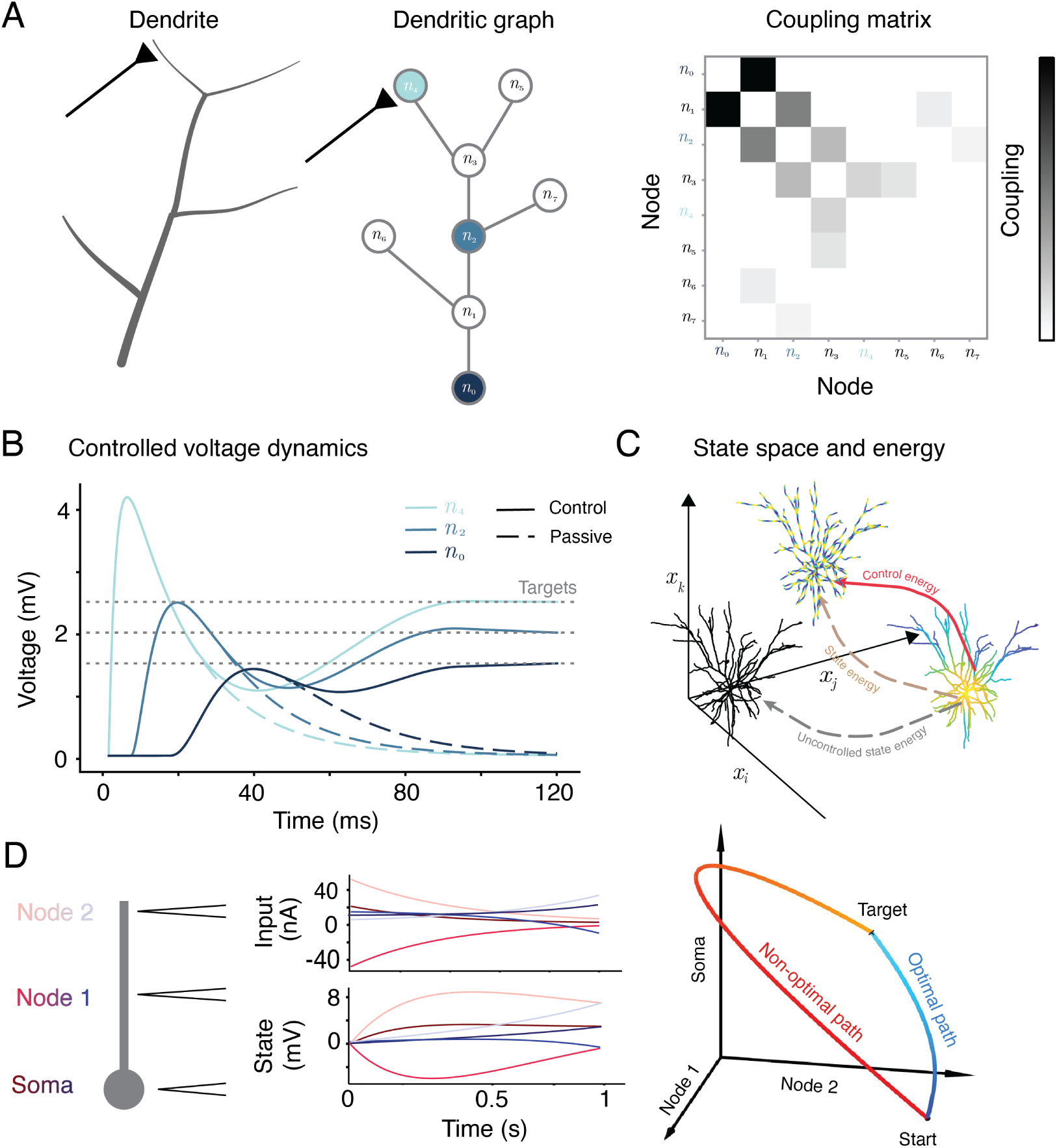
Network control theory applied to dendrites. **A** Left to right: A dendrite receiving synaptic input, the dendrite represented as a tree-structured graph with interconnected nodes, and the coupling matrix between dendritic nodes (excluding self-connections). **B** Evolution of dendritic voltages at different locations due to synaptic (passive, dashed lines) and synaptic plus control (control, solid lines) input. Synaptic input arrives at node *n*_4_ at time 0. **C** Schematic of the evolution of dendritic state under intrinsic dynamics (dashed line) and control inputs (solid lines). **D** Left: a simple example of a soma and two dendritic nodes that receive control inputs. Center: Input current (top) and voltage state (bottom) for optimal (blue) and non-optimal (red) control inputs. The objective was to reach 3.0 *mV*, −1.0 *mV*, and 7.0 *mV* for the soma, node 1, and node 2, respectively, at *t* = 1 s. Right: State trajectories in 3D space for the two cases.

Consistent with this view, pioneering work by Poirazi and Mel showed that complex dendritic integration allows a single cell to perform the computations of a two-layer neural network (Poirazi et al., 2003b), underscoring the importance of dendritic nonlinearities (Cazé et al., 2013; Tran-Van-Minh et al., 2015). More recently, it has also been shown that the complexity of input-output computations in a single neuron with dendrites and dendritic nonlinearities (such as NMDA spikes) is equivalent to that of a deep multi-layer artificial neural network (Beniaguev et al., 2021, 2022). Individual neurons, therefore, behave as high-dimensional complex dynamical systems in their own right. This framework raises the possibility that control theory approaches, which have been successfully applied to neuronal networks (Srivastava et al., 2022; Tang and Bassett, 2018), may also be useful for understanding how the properties of dendritic trees allow them to be driven to functionally useful target states.

## Network control theory applied to brain networks

Control theory is a framework in mathematics and engineering that, when applied to an existing system, helps reveal the inputs that can efficiently shift a dynamical system from an initial state to a desired state in a restricted time interval and, in some cases, track or maintain that target state (Glad and Ljung, 2018).

Network control theory (NCT) is an application of control theory to complex networks where many systems and nodes interact with each other (Cornelius et al., 2013; Liu et al., 2011; Lombardi and Hörnquist, 2007; Pasqualetti et al., 2014), such as power grids (Jawad and Gou, 2013; Liu et al., 2022), epidemic networks (Pastor-Satorras et al., 2015), metabolic networks (Kim and Motter, 2009; Liu et al., 2011), logistic or even social networks (Cremonini and Casamassima, 2017; Dolgui et al., 2018). When applied to high-dimensional complex networks, control theory helps to identify the network inputs (or, more generally, network perturbations) that will drive the network from its current state to a target state (Tang and Bassett, 2018). This allows a system to quickly and efficiently fulfill its desired, or evolved, objectives and can prevent catastrophic dynamics.

In neuroscience, NCT has previously been applied to networks of neurons in C. elegans (Liu et al., 2011; Yan et al., 2017) and more recently to networks of brain regions (Srivastava et al., 2022). In this framework, any state of a network of *n* active neural elements (typically brain regions, but can also be local circuits or neurons) can be formalized as a vector of *n* individual dynamical states **x** = (*x*_1_*, x*_2_*, …, x_n_*) that interact with each other through some function *f* (.), usually modeled as the following, given external control input **u**(*t*):

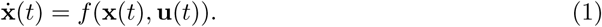

In biological systems, *f* (.) is usually nonlinear. However, many useful results in classic control theory are applicable only to linear systems with additive inputs; hence, a locally linearized version of *f* (.) is often used. NCT allows one to find one or more control inputs (i.e., perturbations: *u*_1_, *u*2*…u_n_*) that can drive the network from an initial state **x**(0) to a target state **x**(*T*) at any desired t=T (Karrer et al., 2020), or shape the network state trajectory in between these points. The trajectory of the network is formalized as a path through the state space in time. It can be described mathematically as a temporal sequence of network states (network state vectors). In this framework, a network state is represented by the physiological activity of all its nodes, i.e., brain regions, at a given time (Karrer et al., 2020; Tang et al., 2020). In brain networks, the control inputs can be internal (endogenous brain activity) or external (brain stimulation). Primarily applied to brain imaging or electrophysiological data, network control theory has offered new insights into the controllability of whole-brain networks under normal, pathological, or externally stimulated conditions. More specifically, NTC has been applied to networks reconstructed from human brain imaging data, showing that the strength of connections between areas determines how difficult or easy it is to reach different brain states (Cornblath et al., 2019; Gu et al., 2015). This framework enabled the discovery of structural network differences that determine how readily the brain can change its cognitive and functional state (Gu et al., 2015). Network controllability can also predict the activation of different brain regions during a working memory task and relate behavioral differences between humans to gender-specific brain rewiring during development (Cornblath et al., 2019). Additionally, Medaglia et al. (2018) found that patients’ performance on the task of appropriate word selection was linked to the controllability of the left inferior frontal gyrus, a brain region strongly associated with language production. A fascinating study has found that low-dimensional dynamics of brain activity align with the spatial patterns of neuromodulatory receptor density, which can alter the effective connectivity between different brain regions and thus the controllability of the brain (Shine et al., 2019).

In pathological brain states, such as seizure dynamics, NCT has provided insights into identifying network regions that control the degree of synchronization among other subnetworks and predicting the spread of seizures before they begin (Khambhati et al., 2016). These controllers can induce pathological states in the brain by reducing the level of push-pull interaction between synchronizing and desynchronizing regions. Another study used control theory to design optimal electrical stimulation protocols for treating patients with epilepsy (Stiso et al., 2019). It has also been shown that the patterns of white matter tracts in the brain allowed for fast and efficient controllability under healthy conditions, but were highly vulnerable to external injuries (Gu et al., 2017).

## A novel proposal: Network control theory applied to dendrites

As argued above, an individual neuron with its dendritic tree can be viewed as a system of interconnected compartments that dynamically pass through different activity states. Biophysical modeling based on cable theory is a powerful approach to simulating dendritic dynamics; graph and cable theory provide two established frameworks to capture the topology (Cuntz et al., 2007) and electrophysiology (Rall, 1959) of dendrites. However, to get a more general idea of the properties of dendritic trees, a broader mathematical theory is needed. Can NCT provide such a complementary formalism to investigate how the spatial and temporal dynamics of neuronal dendrites arise from their underlying biophysics and topology (graph structure)?

Applying NCT to neuronal dendrites depends upon first constructing a graph model in which dendritic nodes are physically linked to each other by edges (Fig. 2A middle). Nodes can be defined in many different ways, for example with uniform length, volume, axial resistance, or using ideas of functional dendritic compartmentalization (Cuntz et al., 2010; Wybo et al., 2019). Having defined the nodes, the underlying graph can be constructed using an electrotonic coupling matrix *A* (Hines, 1993; Cuntz et al., 2007). Second, if a dendrite is a tree of nodes connected by edges, then the state of the dendrite at time *t* can be represented as the pattern of voltages across all nodes of the tree, giving a state vector **x**(*t*). Similar state space formalizations of a dendritic tree have been used before (Rall, 1964; Pakman et al., 2014; Bird and Cuntz, 2016).

Analogously to network state trajectories (Gu et al., 2015; Karrer et al., 2020), the dendrite state trajectories can then be formalized as the temporal sequence of voltage states that the dendrite, as a passive dynamical system (without external inputs **u**), goes through (Fig. 3B).

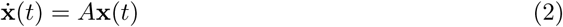

**Figure 3:**
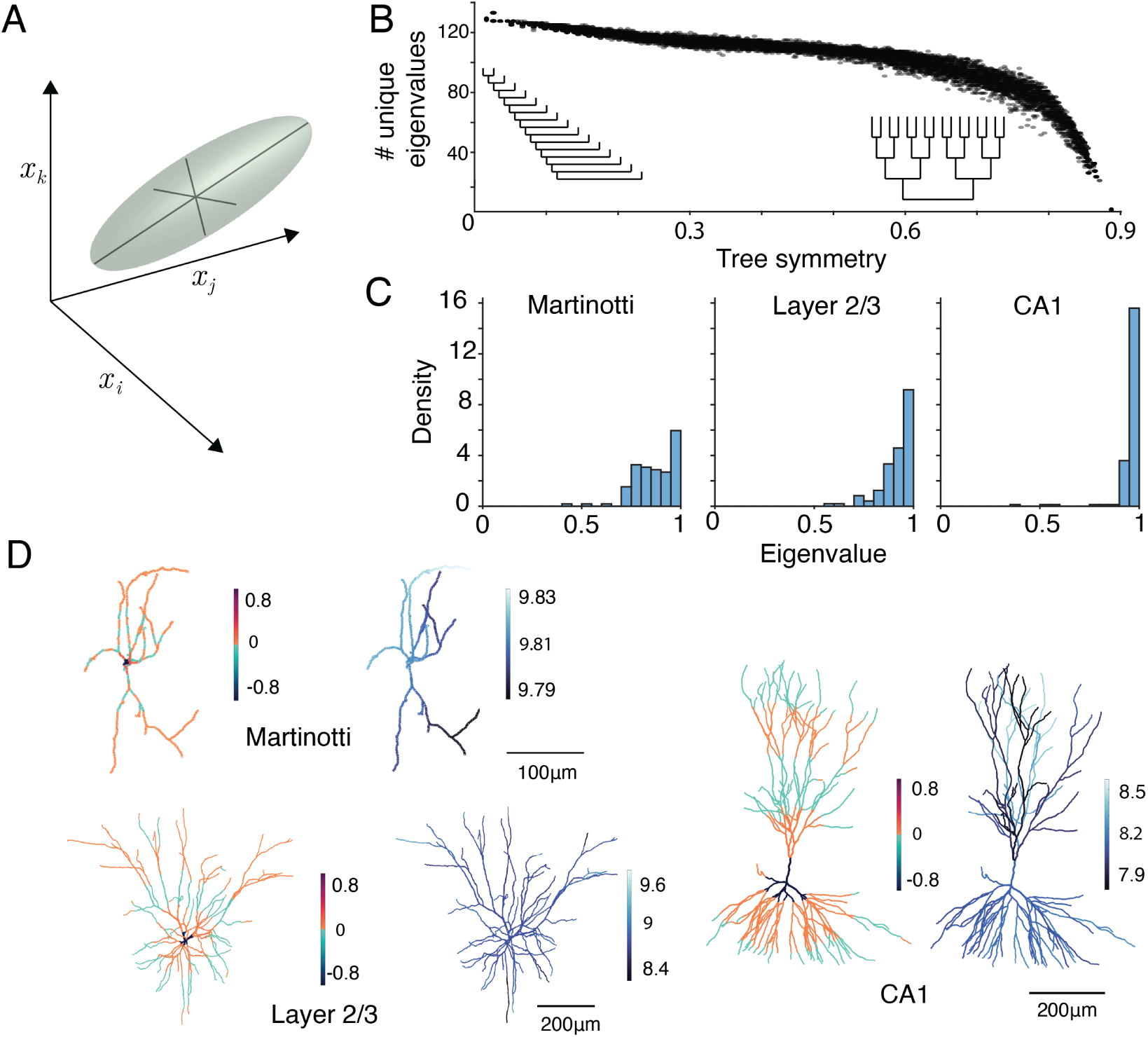
Eigenvalues and eigenvectors of the Gramian matrix. **A** Schematic of eigenvectors in state space. Large magnitude vectors corresponding to large eigenvalues indicate directions that are easy to control. **B** Relationship between the asymmetry of binary trees and the number of unique Gramian eigenvalues. Insets show perfectly symmetric and asymmetric topologies. **C** Distributions of eigenvalues of the Gramian matrix for three neurons, from left to right: a Martinotti interneuron, a cortical layer 2/3 pyramidal neuron, and a hippocampal CA1 pyramidal cell. **D**Eigenvectors corresponding to the smallest (left) and largest (right) eigenvalues of the Gramian matrix mapped onto the above morphologies.

The matrix *A* is the symmetric coupling matrix that is inferred after either linearizing the nonlinear interactions *f* (.) or directly from the passive relationships between nodes in a dendrite. It is an *n*-dimensional matrix, with *n* being the number of nodes in the dendritic graph. This coupling matrix represents the instantaneous transfer resistance between dendritic nodes (Fig. 2A, right), with membrane leak conductances on the leading diagonal. Typically, a stable matrix *A* imposes relaxation of dendritic voltages to rest in the absence of external input (Fig. 2B dashed versus solid lines). The tree topology and symmetric structure of the dendritic coupling matrix *A* impose some unique constraints on the dynamics and controllability of dendrites that we discuss in detail in future sections.

## Controllability of the linearized dendrite

Having established the formal descriptions of the dendritic graph and its dynamical states, we can now use NCT to define the controllability of the dendrite. The usefulness of the controllability concept lies in determining how the dendrite can be directed to particular target states by external or internal control inputs (Gu et al., 2015; Kalman et al., 1963; Karrer et al., 2020; Rahmani et al., 2009). In real neuronal dendrites, such an internal control input can be represented, for example, by excitatory or inhibitory synaptic inputs or other (e.g., neuromodulatory) inputs. An external control input could be local electrical or optogenetic stimulation, which can be applied to the distal or proximal dendrites or to the soma or axon.

According to control theory, a dynamic system - in our case, a single neuron with complete dendritic morphology - is fully controllable if a set of control signals can “steer the internal state of the system from any initial state to an arbitrary accessible final state in a given time interval” (Frank, 2018). Following the formalism of NCT (Karrer et al., 2020; Tang and Bassett, 2018), to account for internal or external input signals injected into selected nodes, we can extend equation (2) by adding a control input matrix *B* multiplied by an injected control input vector **u**(*t*). This is the form of Eq. 1 that is widely used in linear control theory.

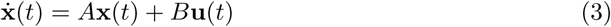

The control input matrix *B* represents the coupling between external input signals to individual dendritic nodes, and has a size of *n*-by-*m*, where *m* is the control input dimension. The input **u**(t) is a control vector, which represents a set of control signals injected into the dendritic tree over time through the subset of *m* control nodes, thereby guiding dendritic activity patterns towards a specific target state. Accordingly, **u**(*t*) is a vector with a size of *m*-by-1 containing *m* control inputs **u**(t) = [*u*_1_(*t*); *u*_2_(*t*); …; *u_m_*(*t*)], applied to all *m* control nodes at each time point *t* (Karrer et al., 2020).

Linear, time-invariant systems are controllable if the following controllability matrix (of size *n*-by-*mn*) inferred from equation (3) has a rank *n* (Kalman et al., 1963)

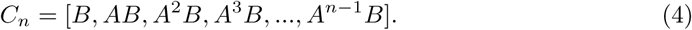

This guarantees that if the rank of *C_n_* is *n*, the dendrite is fully controllable, meaning that the entire dendrite (together with the soma) can achieve any possible state within a finite time *T*. The rank of the controllability matrix determines how many independent vectors can be accessed through the control input (Kim and Bassett, 2020). A set of all such target dendritic states that can be reached from any initial state (using a suitable control input) is called the “controllable subspace” of the system given the input structure.

To illustrate the concept of controllability in a simple neuron example, we considered a ball-and-stick model, representing a soma with two dendritic nodes, leading to a three-dimensional system (Fig. 2D). The objective of this toy example was to drive the soma, dendritic node 1, and node 2 to 3.0*mV*, −1.0*mV*, and 7.0*mV*, respectively, at 1.0 seconds from the start with a zero initial condition. This distribution of voltages is non-monotonic as a function of path length and would be challenging to achieve without techniques from DCT. The connectivity matrix between nodes (A) had −0.7775 *×* 10*^−^*^9^*s^−^*^1^ and −1.5551 *×* 10*^−^*^9^*s^−^*^1^ for the self-coupling values of the soma and dendritic nodes, respectively. The coupling value between the nodes was 0.7775 *×* 10*^−^*^9^*s^−^*^1^. The input matrix was considered to be a unity matrix; i.e., all nodes received external drive with identical resistance values. The controllability matrix (Eq. (4)) for this example was full-rank. As shown in Figure 2D (right, red trajectory), it was possible to design an input signal *u*(*t*) to achieve the target values at *t* = *T_f_* = 1s. Such a generic control input is obtained from (Chen, 1984)

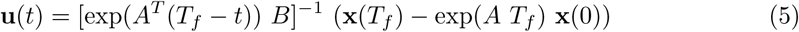

This input design, however, was not optimal; i.e., it did not minimize the input and state signal energies. In the following section, we will provide a framework for designing an optimal control input.

## Energy costs of dendrite controllability: Minimum and optimum control energy

In most engineering and biological systems, it is essential to keep energy costs minimal. Optimal control theory, a branch of control theory, provides a useful formalization of energetic efficiency (Pasqualetti et al., 2014).

Generally, three conditions need to be specified to formulate an optimal control problem (Kirk, 2004): First, the process under control has to be modeled mathematically. For dendritic dynamics, the mathematical description consists of the compartmental model and the state transition equation (3). Second, the physical constraints for the values of the states and the control inputs need to be identified. For example, the weights of synaptic inputs are bounded by their known upper limits of a few nanosiemens. In addition, the ranges of potential voltages as well as calcium concentrations are experimentally well known as physical constraints, excluding physically impossible states of the dendrite. Third, performance criteria or measures need to be defined.

Having been employed in NCT (He et al., 2022; Tang and Bassett, 2018), optimal control theory could also be helpful in estimating the energetic costs of controlling dendrite dynamics. When using optimal control theory for dendrites, the performance can be specified based on their known tasks and computations. This can be, for example, coincidence detection (Remme et al., 2018), pattern separation (Bird et al., 2024), or even more basic objectives such as current transfer (Bird and Cuntz, 2016).

In the mathematical definition of energy, input control energy can be calculated as a squared function of the control input **u**(*t*) required to drive the dendrite from one state to another through a trajectory of intermediate states:

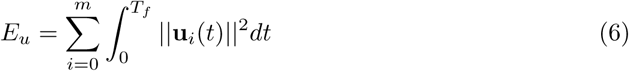

Similarly, the state energy of the system is the sum over the squared states of the system from the initial condition to the target state **x**(T), as follows:

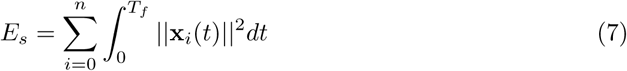

which reflects the length of the transition trajectory from *t* = 0 to *t* = *T_f_*.

Optimal control theory provides a framework for minimizing not only the control energy *E_u_*, but also the state energy *E_s_* of a controllable system. Whereas optimal control energy (OCE) considers both the control input energy and the length of the state transition trajectory, minimal control energy (MCE) includes only the required input energy costs *E_u_* regardless of the state trajectory (Karrer et al., 2020).

The optimal control input is determined by the minimum of the cost function J (Kirk, 2004):

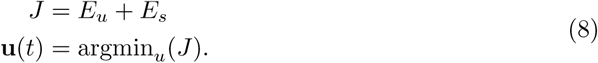

It can be shown mathematically that the optimal input signal for a given pair of *A* and *B* in equation (3) follows:

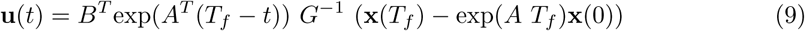

where *G* is the Gramian matrix of the system defined by *A* and *B* as follows:

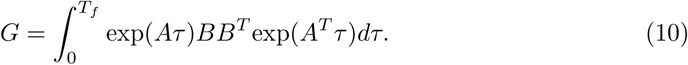

Although the energy terms defined in equations (6) and (7) follow mathematical definitions for energy contents of signals, it will be interesting to compare these measures with standard energy cost measures based on ion counting and inferred ATP costs (Jedlicka et al., 2022; Remme et al., 2018) from compartmental modeling of neuronal dendrites.

To illustrate the advantage of optimal control theory, we applied the optimal control design following (9) to the example in Figure 2D, aiming to achieve the same target values. This framework resulted in state and input trajectories that required less signal energy (blue line) compared to the non-optimal input, despite sharing the same initial and target states.

The Gramian matrix in (10), essential for calculating the control input for minimal state and input energy, is always symmetric and has non-negative entries, and hence has real positive eigenvalues for a controllable system. There is an inverse relationship between the magnitude of a Gramian eigenvalue and the energy required to control the system along its corresponding eigenvector (Chen, 1984). Therefore, a target value aligned with the eigenvector corresponding to the largest eigenvalue of the Gramian requires the least input energy, and a target value aligned with the eigenvector associated with the smallest eigenvalue is the most costly desired point to achieve (Fig. 3A).

Another interesting property of the Gramian originates from the Perron–Frobenius theorem (Perron, 1907; Frobenius, 1912), which states that for any non-negative matrix, the largest eigenvalue is unique and positive and its corresponding eigenvector has all positive (or all negative) values. This immediately implies that the minimum control input would either excite all nodes or inhibit all nodes, but it cannot suppress a subset of states in favor of elevating other states. This indicates that it would be easiest to either excite the dendrite and soma together or to suppress all nodes.

## Unique features of dendrite control theory

The tree structure of dendrites gives rise to some unique features that distinguish DCT from broader NCT. In particular, the coupling matrix between passive dendritic nodes is real-valued and symmetric, which implies the following:

### No intrinsic oscillations

Basic theorems in linear algebra have proven that in symmetric matrices, all eigenvalues are real (Strang, 2016). This implies that in passive dendrites, no oscillations arise due to the intrinsic dynamics. Hence, all oscillatory dynamics need to be provided by the input signal. For active dendrites, nonlinearities might break the symmetry of the linearized matrices *A* and so allow intrinsic oscillations. This aligns with predictions from classical cable theory (Rall, 1964; Rall and Rinzel, 1973) and highlights the necessity of active properties for computationally useful sustained local oscillations (Lengyel et al., 2005; Remme et al., 2009).

### ‘Complex’ dendrites have less diverse energy demands for control

Different tree structures can give rise to different eigenvalue distributions for the underlying connectivity matrix, and hence different eigenvalue distributions for the Gramian matrix. In graph theory, it is known that star-like graphs have an eigenvalue spectrum mostly close to zero, with an outlier eigenvalue with a very large value. However, cable-like graphs have a more uniform distribution of eigenvalues (Braga et al., 2013; Nakatsukasa et al., 2013; Rohleder, 2017). For uniform binary trees with different asymmetries (Liao et al., 2023; Uylings and van Pelt, 2002), we also found that more symmetric trees have fewer unique eigenvalues (Fig. 3B). More distinct eigenvalues imply greater variability in the control energies required to reach different states; cable-like and asymmetric neurons are likely to have a strongly preferred control direction, whereas symmetric and star-like neurons can have more diversity in response properties.

For real cells with different morphologies, i.e., a Martinotti interneuron, a cortical layer 2/3 pyramidal cell, and a hippocampal CA1 pyramidal cell, we found that eigenvalues of the Gramian matrix became more concentrated as complexity increased (Fig. 3C). When considering the eigenvectors associated with the smallest and largest eigenvalues of the Gramian matrix, we were able to identify the directions in which it was easiest and hardest to drive a neuron. For all three examples, the hardest control input was that which isolated the soma from the rest of the dendritic tree (Fig. 3D, left morphologies) and drove it in an opposite direction to the proximal dendrites. The picture for the easiest control input was more complex (Fig. 3D, right morphologies). All neurons were easiest to drive in a consistent direction (as a result of the Perron-Frobenius theorem). The magnitudes of the eigenvector elements remained very close together, with complexity leading to slightly more diversity between elements. This reinforces the idea that complex cells, such as star-like neurons or those with many branch points, are relatively easier to drive to more diverse dendritic states.

### Peripheral inputs are sufficient for controllability

More recent theories on control theory applied to systems with tree-structured connectivity matrices have shown that applying control inputs to all peripheral nodes is sufficient to achieve full system controllability (Chao and Ji, 2021; Lin, 1974; Hsu and Yang, 2019). To check this in our simple three-dimensional system of a soma and two dendritic nodes in Figure 2D, we provided external control only to the last distal node (node number 2) and aimed at achieving a target state of 3.0 *mV*, −1.5 *mV*, and 2.0 *mV* at the soma, node 1, and node 2, respectively. We used the optimal control design in eq (9), and showed that the resulting input current can indeed achieve the desired target state. Next, we studied the application of this theory in more realistic higher-dimensional dendritic morphologies and analyzed their controllability under input pruning.

A relevant dendritic measure for the proximity of a node to the periphery or the soma is the Strahler order. Strahler order is defined to be equal to one for peripheral branches, and increases by one every time branches with equal Strahler order meet towards the soma (Strahler, 1957; Vormberg et al., 2017). To examine the robustness of the controllability of the neuron (all dendritic compartments and the soma) in the presence of input pruning, we performed two computational experiments: 1) We removed inputs to nodes with the smallest Strahler orders first and continued pruning with increasing Strahler order; 2) We removed inputs with descending Strahler order. Our measure of the system’s controllability was the rank of the Gramian matrix for (*A, B*) (Eq. 10), with *B* losing its rank progressively as a result of input pruning. In both cases, the total number of control input dimensions decreased on average. However, we observed an interesting pattern in the resilience to input pruning: the neuron maintained its full controllability when the inputs were removed with decreasing Strahler order, i.e., the scenario in which the control inputs to the dendritic compartments in the periphery were kept until the late stages of the experiment rendered a more robust controllability than that where peripheral inputs were removed first (blue versus orange in Fig. 4A).

**Figure 4:**
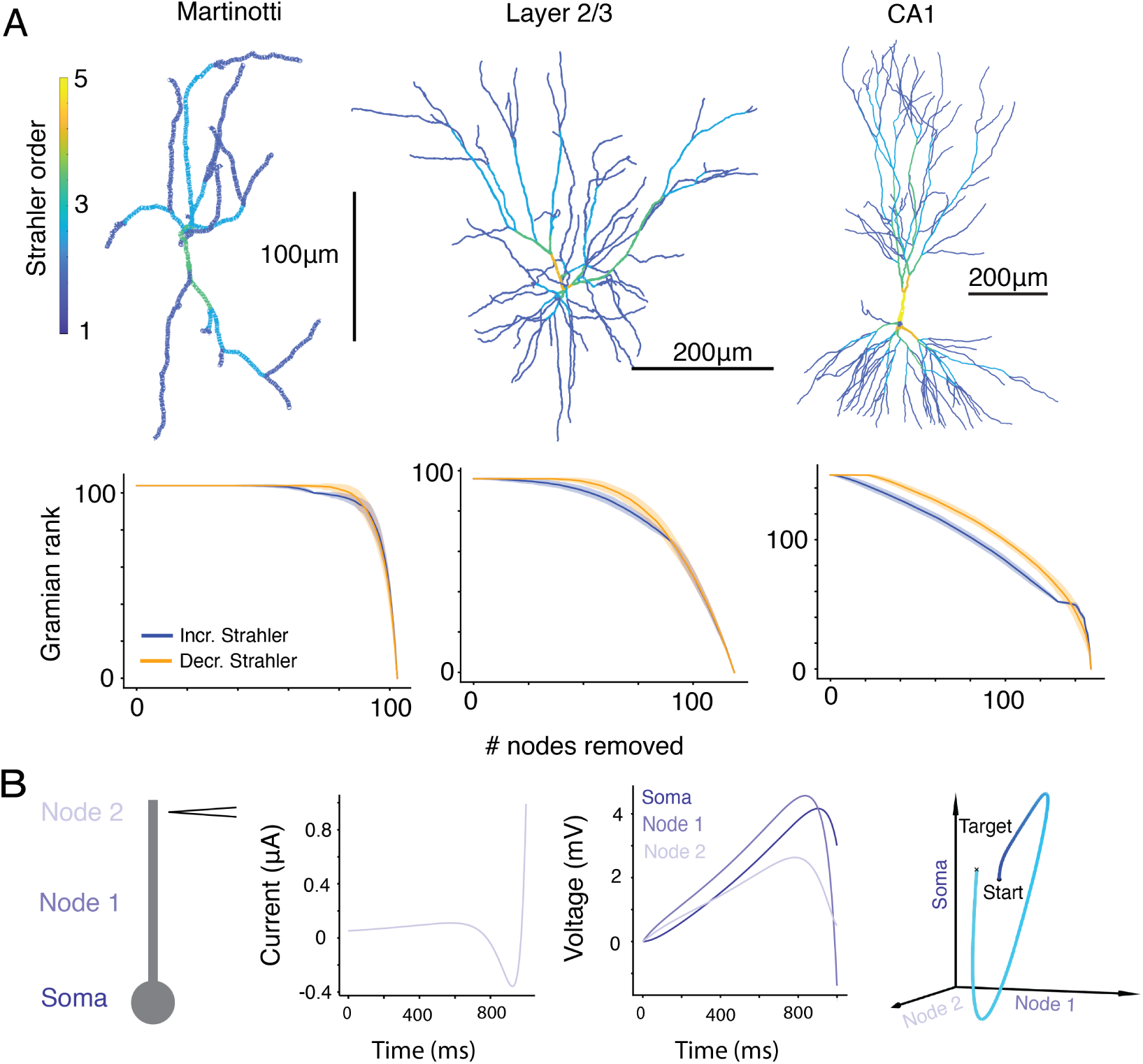
Peripheral inputs are sufficient for control. **A** Top row, left-to-right: morphologies of a cortical Martinotti interneuron, a cortical layer 2/3 pyramidal neuron, and a hippocampal CA1 pyramidal cell, colored by Strahler order. Bottom row: Rank of Gramian matrices for the neurons when inputs are removed peripherally first (blue), and proximally first (orange). **B** Three-node dendrite (first left), with control input (second left) only in the peripheral node, can achieve any desired state at *t* = 1s (second and first right).

Somatostatin (Sst) interneuron subtypes are a prominent example of presynaptic cells that preferentially target peripheral dendrites and could provide such robust control signals (Silberberg & Markram, 2007; Petilla Interneuron Nomenclature Group, 2008; Gidon and Segev, 2012). Our results hence suggest that interneurons such as Sst cells could provide full controllability of the entire dendrite, while perisomatic interneuron subtypes such as Parvalbumin cells provide local, or output, controllability, mainly targeting the somatic state. The controllability granted by dendritic tree structure also implies a surprising robustness to changes in specific inhibitory locations as long as general peripheral targeting remains. In fact, experimental evidence suggests that inhibitory input synapse locations are highly dynamic, particularly those that innervate peripheral spines Boivin & Nedivi. (2018). This potentially allows continuous fine-tuning of inhibition whilst maintaining controllability. For instance, Sst cells have also been implicated in controlling the plateau state of the dendrite (Vaasjo et al., 2026). Theoretical work has also shown how Sst inputs can alter local input resistance (Gidon and Segev, 2012), and therefore change patterns of local excitability across a dendrite (He et al., 2026). DCT shows that peripheral targeting allows these functions to be carried out robustly.

Another interesting theoretical observation was that dendrites with fewer branch points required fewer inputs to be fully controllable. According to control theory, cable-line trees require at least one input in an asymmetric location to achieve full controllability (Chapman and Mesbahi, 2013; Hsu and Yang, 2019). The typically simpler morphologies of inhibitory neuron dendrites, such as Martinotti cells, imply that these cells require fewer control inputs and would therefore be more resilient to input turnover or loss of synaptic inputs (Keck et al., 2011). In other words, these cells can maintain their desired state even under strong structural disruptions and continue to exert inhibitory control over network dynamics.

### Dendrite dimensionality can be adaptive

Dendritic trees *in vivo* are dynamic, making the coupling matrix *A* state-dependent and allowing adaptive gating of control signals. For instance, mechanisms such as neuromodulation can decouple the apical tuft of layer V pyramidal cells from the soma, with potential consequences including the (temporary) loss of consciousness (Larkum et al., 2009). Even shunt from excitatory and inhibitory synaptic inputs can change the effective compartmentalization of a neuron (Cuntz et al., 2010; Wybo et al., 2019), altering the number of functionally relevant voltage states a neuron can take and therefore the dimensionality of *A*. It is possible that higher-level systems may alter the dimensionality of dendrites, and thus their control properties, leading to a hierarchy of control in the brain (Pezzulo and Cisek, 2016).

### Average dendrite controllability via single nodes is poor

Average controllability for each node of the system is a metric that refers to the average required energy to drive the whole system to any desired state by providing control input only to that node. We have shown previously that with an input dimension equal to one, the rank of the Gramian matrix decreases significantly, rendering the system uncontrollable. Therefore, we conclude that dendrites have poor average controllability through single nodes.

## Potential applications of dendrite control theory

### Synaptic plasticity

Many open questions in the field of dendritic dynamics remain to be addressed, and the framework of DCT offers a promising new path forward. DCT could provide insights into how passive and active dendritic mechanisms, in concert with excitatory and inhibitory synaptic inputs as well as neuromodulatory signals, jointly constrain and optimally control both local and global dendritic states. These states include functionally significant events such as dendritic plateau potentials, dendritic spikes, bursts, and BAC firing (Bittner et al., 2017).

Importantly, this control perspective can also be extended to dendritic, synaptic, and intrinsic plasticity. Neuromodulatory influences on synaptic plasticity - such as those formalized in three-factor learning rules (Frémaux and Gerstner, 2016) - can be naturally interpreted as a form of control. DCT and its control energy formalism (see above the section with equations (6), (7), (8), (9)) are particularly well suited to address questions related to the hypothesis that neuronal dendrites improve neural network performance while minimizing energy costs (Yu and Yu, 2017; Jedlicka et al., 2022; Makarov et al., 2023), potentially ensuring efficient coding of information (Seenivasan and Narayanan, 2022). As mentioned above, DCT offers a natural mathematical language to study how plasticity mechanisms shape the fundamental trade-offs between dendritic function, flexibility, robustness, and the energy-efficiency of neural networks (Jedlicka et al., 2022).

### Dendritic morphology

The morphology of dendrites appears to be constrained by an optimal balance between maximizing the speed of signal propagation (i.e., minimizing the path length to the soma) and minimizing the total cable length (Cuntz et al., 2007, 2010). This trade-off strongly influences the number of dendritic branches for a given neuron type. However, the number and topological organization of dendritic branches are also relevant for the efficiency of control inputs, both through defining ‘easy’ and ‘hard’ state directions via eigenvalue structure and through allowing different numbers of control input locations. There appears to be a trade-off here between highly branched dendrites, which can easily take diverse dendritic voltage states, and simpler dendrites that require fewer synapses to control, hence improving the resilience to structural disruptions. Incorporating topological considerations from DCT alongside geometric optimality could lead to new insights into the morphologies of different neuron types with different functional roles.

### Synaptic placement

A potential application of the DCT is, given a fixed morphology, the identification of optimal synaptic parameters, including placement, weights, and kinetics, for both excitatory (Magee, 2000) and inhibitory inputs (Gidon and Segev, 2012) to drive a neuron to these states. A recent study (He et al., 2026) applied a similar framework to search for general principles in inhibitory targeting, finding a complex relationship between dendritic excitability, excitatory inputs, and effective inhibitory sites.

Previously, we have shown how tree depth and symmetry can influence controllability; here, we explored how different spatial distributions of control inputs lead to different energy requirements in the formation and maintenance of dendritic plateau potentials. We considered a hippocampal CA1 pyramidal cell model, discretized to 149 dendritic compartments, which, together with the soma, comprised a 150-dimensional tree model. The aim was to design external control signals that lead the entire dendrite from a randomly distributed initial condition to three distinct plateau values in a 50ms time interval, and maintain that state for a further 150ms. To examine the effect of input dimensionality (*m*) on the energy required to accomplish such a task, we provided external input to all peripheral nodes (113 out of 150 nodes) in addition to subsets of the remaining nodes, in increments of 5, until all nodes received inputs.

Our results (Fig. 5) indicate that optimal inputs require the least control energy to achieve a dendritic plateau state if “all” nodes of the dendrite and soma receive input signals (*m* = *n* = 150 in equation (9)). In other words, restricting inputs to fewer nodes (*m < n*), progressively increases the energy demand to achieve the plateau values within a given time window. This could explain why, among many other reasons, evolution has favored multiple globally distributed inputs for their energetic efficiency (Kasthuri et al., 2015; Motta et al., 2019).

**Figure 5:**
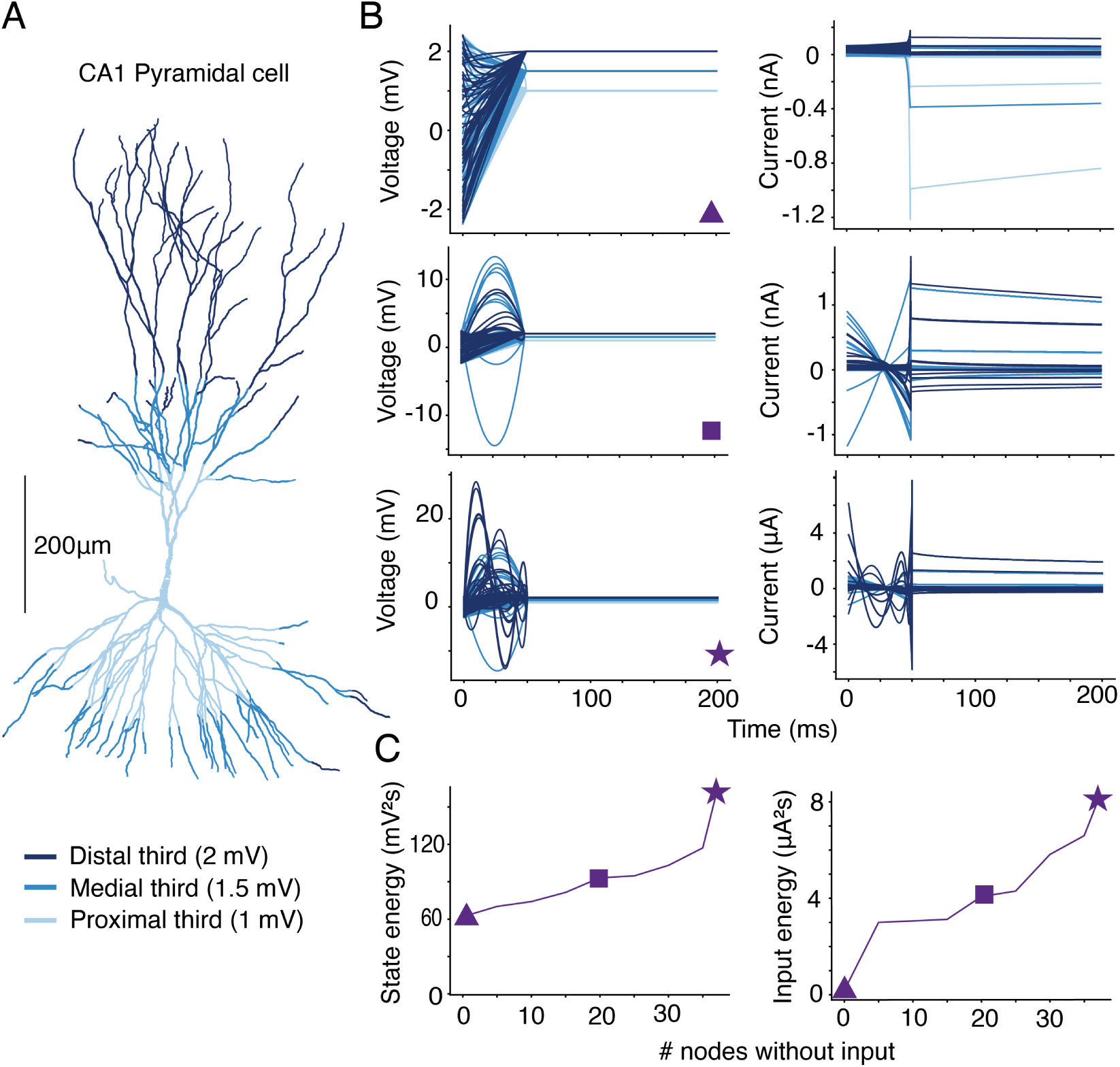
Control energy is lower for distributed inputs. **A** Morphology of a hippocampal CA1 pyramidal neuron, colored by distance from the soma. **B** State (left) and control input (right) when all nodes can receive input. **C** Minimum state (left) and control (right) energies required as a function of the number of inputs removed.

### Dendritic computation

Dendrites are involved in the implementation of complex computations, such as coincidence detection, feature binding, and motion detection (London and Häusser, 2005; Remme et al., 2018; Tran-Van-Minh et al., 2015). The eigenvalue spectrum of a neuron’s Gramian matrix provides a way to evaluate which dendritic states are favored from an energy point of view. By analyzing the preferred directions emerging from the eigenvectors of the Gramian matrix at different timescales *T*, we can infer how well neurons might perform particular computational functions for different sets of synaptic inputs (Cazé et al., 2017; Tran-Van-Minh et al., 2015).

Many network-level computations, such as sequence generation and motor function, rely on the order or timing of spikes in individual neurons (Srivastava et al., 2017; Ortega et al., 2019; Egger et al., 2020; Trusel et al., 2026; Shapcott and Bird, 2021; Yiling et al., 2023) For example, in the Higher Vocal Center (HVC) of songbirds, the timing of the neuronal activity is critical for stereotypic song generation (Hahnloser et al., 2002; Long et al., 2010; Trusel et al., 2026). Grasping behaviors in primates are also coordinated by sequences of spikes in the motor cortex (Kalaska et al., 1997). DCT allows the precise determination of the inputs necessary to drive a neuron and its dendrites to threshold with any arbitrary delay.

### How can DCT guide experimental research?

A useful application of DCT may be the identification of optimal light-based (optogenetic) control strategies for the experimental manipulation of dendritic neuronal dynamics. It has been suggested (Grosenick et al., 2015) that mathematical approaches in control theory (Huggins and Paninski, 2012; Pakman et al., 2014; Paninski, 2010; Pnevmatikakis et al., 2012) can be combined with light-based stimulation and fast calcium and voltage imaging of dendrites (Grienberger et al., 2015; Park et al., 2024). For example, 3D two-photon holographic uncaging has recently been used to explore the role of dendritic sodium spikes and plateau potentials in the discrimination of multibranch synaptic input patterns in L5 pyramidal neurons (Xiao et al., 2024). These optical and mathematical methods can be used to reveal the naturally preferred directions of activity for a dendrite, using the eigenvectors of the Gramian matrix. An interesting test for experimentalists would be to see which regions of different cell types are relatively easy to excite: Is the soma favored relative to the distal parts, or can distal regions be more easily excited?

Studying difficult-to-control directions may also help with better understanding the role of different morphologies in neuronal functions. In our examples, we showed that the most difficult-to-control state involves designing an input to drive the membrane potential for the soma (and nodes in its proximity) in the opposite direction compared to the rest of the dendritic nodes. Experimental design could test the subspaces of neuronal states that can be controlled for neurons with different morphologies and the energy required to drive them in such non-preferred directions (Kao and Hennequin, 2019). For example, how easily can the somata of excitatory and inhibitory neurons with different morphologies be isolated from the rest of their dendritic trees? The answer to this question identifies whether certain cell types can selectively be driven into responsive or non-responsive states through top-down control from other brain regions. For example, Vasoactive Intestinal Peptide (VIP) interneurons in the visual cortex during attentive states or upon observing novel objects are known to express elevated activity due to top-down input from the frontal cortex (Bastos et al., 2023; Zhang et al., 2014; Garrett et al., 2020). These cells also receive excessive inhibitory drive from Sst cells and usually exhibit low firing rates. Given the bipolar dendritic morphology of VIP neurons, would it be easier to drive them into excitable states compared to other inhibitory subtypes when the rest of the dendrite receives dominant inhibitory drive? The answer to this question can relate the dendritic morphology to specific behavioral and functional states required for a specific task.

Another prediction arising from our DCT formalism is that dendrites with a less branched structure, such as those found in inhibitory interneurons, are more resilient to pruning of synaptic inputs and can remain controllable under severe input disruption (Fig. 4). As inhibitory subtypes play a significant role in maintaining the function and stability of brain networks, it would be interesting to perturb the synaptic inputs to different excitatory and inhibitory neuronal subtypes and examine their responsiveness at the network level and compare their resilience to the control condition. For example, to examine the resilience of the sequential pattern of the High Vocal Center (HVC) of birdsong to disruptions in synaptic input, it would be interesting to study the effect of AMPA and Glutamate receptor blocking on the timing of activity for excitatory and inhibitory subtypes during singing.

## Limitations and challenges: Control theory for nonlinear, plastic, stochastic, feedback-regulated, and synapse-driven dendrites

The concepts of controllability and optimality mentioned above are based on linear control theory for time-invariant and deterministic systems. However, dendrites have a number of important features that are not well-modeled by systems of this type. Here, we highlight some of the challenges.

### Nonlinearities

Dendrites display a range of nonlinear properties that are essential to their computational and functional relevance (Cazé et al., 2013; Dembrow and Spain, 2022; London and Häusser, 2005; Poirazi et al., 2003a; Tran-Van-Minh et al., 2015). The existence of active dendritic compartments can cause dendritic state trajectories to split and separate rapidly; whether a local voltage crosses an action potential threshold or narrowly misses it will drastically change subsequent states. This makes the matrix *A* in Eq. (3) state-dependent. A further important nonlinear effect is caused by saturation of conductance-based synapses (Tran-Van-Minh et al., 2015; Cazé et al., 2017): when local voltages approach the synaptic reversal potential, future excitatory inputs will be integrated sublinearly. This makes the matrix *B* in Eq. (3) state-dependent.

While designing control for nonlinear systems remains challenging, novel control approaches are being developed for nonlinear biological systems (Atay et al., 2026; Cornelius et al., 2013; Díaz-Seoane et al., 2023; Lopez and Slotine, 2021; Morrison et al., 2021; Russo and Slotine, 2011; Salfenmoser and Obermayer, 2022, 2023; Whalen et al., 2015; Zañudo et al., 2017), which could also be applicable to nonlinear dendrites to deal with this challenge. For example, the advent of new machine learning tools facilitates the application of data-driven control theory to highly nonlinear systems (Brunton and Kutz, 2019; Shakeri and Mesbahi, 2025). In some cases, linear control models have served as useful approximations of nonlinear brain networks (He et al., 2022) and neuronal dynamics (Bicknell and Latham, 2024; He et al., 2026; Hennequin et al., 2014; Kao and Hennequin, 2019; Moore et al., 2024; Sani et al., 2021). More rigorously, the nonlinear dynamics of the dendrite can be adaptively linearized around specific states (Schöll et al., 2016; Slotine and Li, 1991). Another promising approach is to use the Koopman operator theory (Brunton and Kutz, 2019; Proctor et al., 2018), which enables a low-dimensional nonlinear system to be represented as a linear system, usually infinite-dimensional. This linearization facilitates the application of control theory in nonlinear systems (Brunton and Kutz, 2019).

### Time dependence and plasticity

Dendrites exhibit significant time variance in their biophysical properties due to intrinsic plasticity (Debanne et al., 2019; Frick and Johnston, 2005; Sjöström et al., 2008). Extrinsic changes, for example, in input synaptic weights, can also change the responsiveness of the dendrite to given inputs.

Time-invariant control theory can be adjusted to its time-varying versions. If we approximate the dendrite as a linear, time-variant system, then *A* and *B* will be time-variant matrices. Intrinsic changes of dendrites (Debanne et al., 2019; Frick and Johnston, 2005; Sjöström et al., 2008) can be incorporated in *A*(*t*), and extrinsic (e.g., synaptic) changes in the control input vector can be incorporated in *B*(*t*) (or **u**(*t*)), and subsequently, DCT could be applied locally in time (He et al., 2026). These approaches usually rely on numerical simulations. In fact, in neural networks, linear control theory has recently been used to account for experimentally well-known forms of synaptic changes, namely spike-timing-dependent plasticity (STDP, Bicknell and Latham (2024); Lagzi (2021, 2024); Moore et al. (2024)). This method could similarly be used to deal with a time-varying dendrite.

### Stochasticity

Both synaptic (vesicle release) and intrinsic (opening and closing of ion channels) properties of dendrites have a stochastic nature (Anwar et al., 2013; Diba et al., 2006; Schneidman et al., 1998). This stochasticity can cause neuronal response variability. While trial-to-trial variability has been observed in the response properties of a wide range of neurons, some neurons, particularly those involved in motor actions, have precise response timings (Hahnloser et al., 2002; Srivastava et al., 2017; Ortega et al., 2019; Egger et al., 2020; Trusel et al., 2026). The biological control mechanisms that can cope with intrinsic and extrinsic sources of stochasticity are not well understood (O’Leary et al., 2014; Yang and Prescott, 2023). Control theory applied to stochastic systems could provide a roadmap to better understand these mechanisms. In fact, linear, time-invariant, and deterministic dendrite models can be expanded by incorporating randomness. Stochastic optimal control approaches have been used to control stochastic neurons (Feng and Tuckwell, 2003; Iolov et al., 2014; Renault et al., 2018) and their networks (Bielievtsov et al., 2016; Todorov, 2005; Wells et al., 2015). These approaches might also be employed in dendritic dynamics. Dendrites could be represented and simulated as dynamical hybrid systems (Ju and Bassett, 2020; Wells et al., 2015) with deterministic and stochastic components that smoothly transition while incorporating abrupt random changes between different state trajectories.

### Feedback

In this article, we used concepts from open-loop control theory, yet dendrites operate in a network with numerous regulatory feedback loops. For instance, dendritic excitability is modulated by inhibitory feedback loops, mediated by dendrite-targeting (e.g., Sst) GABAergic interneurons, which influence the likelihood of future synaptic dendritic integration (O’Leary et al., 2014; Yang and Prescott, 2023). Moreover, neuromodulatory systems such as dopamine or acetylcholine pathways provide top-down feedback that dynamically tunes dendritic responsiveness based on functional or computational context.

Feedback control methods have already been studied at the level of behavior (Cowan et al., 2014; Humbert et al., 2024), networks (Batista et al., 2010; Bielievtsov et al., 2016), but also single neurons. For example, control theory has been proposed to account for the homeostasis of neuronal excitability in the context of intrinsic plasticity that is linked to adaptive (global and spatially local) changes in ion channels (Drion et al., 2015; O’Leary, 2018; O’Leary and Wyllie, 2011), and also to model synaptic plasticity based on dendritic error coding (Meulemans et al., 2022; Senn et al., 2023). In general, feedback control is very efficient in suppressing 1) internal or intrinsic disturbances in the system, 2) instabilities of the system caused by changing dynamics of interaction between components, and 3) stochastic external disturbances (Brunton and Kutz, 2019). This might shed light on why biological neurons with their dendrites use many feedback mechanisms that could be conceptualized within the DCT.

### Synaptic input as control input

The classical theorems of control theory rely on continuous, time-varying inputs. This could be achieved by an experimentalist injecting current into a dendrite. Biologically, however, input to a neuron is dominated by chemical synapses. These are discrete events in time and have finite size.

The problem of limited input options is covered with the theories of quantized control (Curry, 1970). These are often implemented in closed-loop control systems, for example, designing optimal optogenetic control inputs based on neural responses (Bolus et al., 2021). For systems without feedback, discrete inputs have been used for epidemic control (Chen et al., 2026) and for shaping neuronal interspike intervals to synchronize oscillations (Dasanayake and Li, 2014). Alongside existing quantized control approaches, mean-field theory could be applied to bridge the gap between discrete inputs and continuous signals. Such approximations are common in theoretical neuroscience due to the discrete nature of spikes (Amit and Tsodyks, 1991; Brunel, 2000; Di Carlo et al., 2026). This could allow DCT to be applied directly to discrete-input systems in some circumstances.

## Summary and outlook

Neurons need to maintain complex states *in vivo*, with voltages potentially varying across the dendritic tree. This allows learning and memory (Ebner et al., 2019; Graupner and Brunel, 2012), and is essential for reliable motor outputs (Hahnloser et al., 2002; Srivastava et al., 2017; Ortega et al., 2019; Egger et al., 2020; Trusel et al., 2026). We have introduced Dendrite Control Theory (DCT) as a novel way to formally analyze how such useful states might be attained and maintained. DCT builds on and extends the view that even a single neuron can be seen as a complex system of interconnected compartments or computational units (Beniaguev et al., 2021; Poirazi et al., 2003b), taking inspiration from the application of NCT to computations, plasticity and the homeostasis of single neurons (Drion et al., 2015; Moore et al., 2024; O’Leary, 2018; O’Leary and Wyllie, 2011; Senn et al., 2023).

We have outlined some of the unique features of DCT that are rooted in the underlying tree structure of the dendritic connectivity matrix. More specifically, we showed that applying control signals to the distal nodes provides a sufficient condition for full dendritic controllability, a function that could be associated with Sst cells, given the peripheral synapses that they establish onto excitatory cells. Such synapses have relatively more dynamic weights (Berry and Nedivi, 2017; Boivin & Nedivi., 2018). This could allow the fine-tuning of control inputs. The resilience in controllability to changes in peripheral inputs also allows control to be maintained at the single-neuron level while circuit-level connectivity changes. Further, we explored how the morphology of the dendrite affects the energy required to control the dendrite along different directions in the state-space. In particular, we found that easy directions for control inputs uniformly excite or inhibit neurons, whereas harder control directions involve isolating the somatic compartment from the dendritic tree.

DCT can provide conceptual and mathematical tools for finding optimal control inputs that can drive the dendrite to diverse functional states. We provided examples of control design to achieve different levels of dendritic plateau states at a specific point in time, for different input dimensions. We showed that restricting the input dimension could give rise to an increase in energy demands: more synapses require less control energy, but require more axonal cable and metabolic maintenance. Another trade-off is between robustness and complexity. Cable-like neurons can be fully controlled with few inputs but have relatively large differences in the control energies to reach different states. Highly branched neurons, on the other hand, require more inputs to fully control, but have smaller differences in the control energies required to reach diverse states. Hence, the DCT framework can help formalize the fundamental trade-offs (Del Giudice and Crespi, 2018; Jedlicka et al., 2022; Makarov et al., 2023; Pallasdies et al., 2021) between functional effectiveness, flexibility, robustness, and energy efficiency of multi-objective dendritic function.

There are, however, challenges to be resolved with DCT. As discussed above, its application to time-variant, nonlinear, stochastic, and feedback-related features of dendritic states has not been explored in this paper. In addition, some aspects of dendritic and, in general, brain dynamics might be beyond formalization and therefore not reducible to mechanistic computation (Jaeger et al., 2024; Potter and Mitchell, 2024). However, despite its current limitations, we suggest that DCT can be a useful tool to understand the requirements to control dendrites of different cell types with different morphologies. Importantly, DCT could be integrated with existing control theory in neuroscience at the level of neurons (Drion et al., 2015; Moore et al., 2024; Senn et al., 2023), neural circuits (Hennequin et al., 2014; Kao and Hennequin, 2019), large brain networks (Gu et al., 2022; Liu et al., 2011) up to cognition (Stocker et al., 2023) and behavior (Humbert et al., 2024; Kessler et al., 2024). Future research can explore the development of a multi-scale control theory in neuroscience (Pezzulo and Cisek, 2016) and study self-organization at multiple scales through the lens of control theory. This could provide insights into the intricate interplay between multiple scales and layers (Nakahira et al., 2021) of the behaving organism.

Predictions from DCT could be compared with experimentally observed properties to better understand the preferred functions and constraints of real neurons (Boivin & Nedivi., 2018; Callan et al., 2021; Kirchner and Gjorgjieva, 2022; Rößler et al., 2023). For example, the energy demands to control different neuronal cell types with diverse morphologies, and their differences in easy-to-control directions, could be tested experimentally. This could provide insight into the computations carried out by different cell types. Moreover, perturbation experiments could identify the robustness of neuronal subtypes to input disruptions and identify the major players in different circuits in the brain.

## Glossary

Average controllability: The ability of a single node to drive the system to any other state.
Control energy: The energy of the control signal (*u*(*t*)) required to drive a dynamic system from its initial state to a desired target.
Controllability: Whether a system can be driven to any possible state by a given set of inputs.
Controllable subspace: The set of all system states that can be driven to an arbitrary target state by applying an external control input.
Dendritic control theory (DCT): The application of network control theory (NCT) to single neurons.
Minimal control energy (MCE): The minimum possible magnitude of control input that can steer a dynamic system from an initial state to a desired target state.
Network control theory (NCT): The application of techniques from control theory to networks.
Optimal control energy (OCE): The magnitude of control input that can steer a dynamic system from an initial state to a desired target state whilst minimizing total control and state energies.

## Acknowledgement

This work was supported by the DFG/ANR project DyndriteSymphony (JE 528/15-1 No. 545518567, CU 217/7-1 No. 62280873, No. 514483642, ANR-24-CE92-0071) to PJ, HC, AG, BG and RC, and Collaborative Research Centre/Transregio 384 (CRC/TRR 384) INCODE, ‘Inhibitory neurons: shaping the cortical code’ (DFG project number 514483642) to AG.

The authors used Grammarly and Google’s AI Overview tool to correct typos and grammar in the text and to improve readability.

## Notes

### Competing Interest Statement

The authors have declared no competing interest.

